# Dye-cycling DNA origami rotors for long-term tracking of transcription at base-pair resolution

**DOI:** 10.64898/2026.01.21.700733

**Authors:** Amanda Wacker, Ryan J. Fantasia, Brian J. Tenner, Jerry Wu, Nicholas A. Monell, Pallav Kosuri

**Affiliations:** Department of Molecular Biology, University of California, San Diego, La Jolla, CA 92092; Salk Institute for Biological Studies, La Jolla, CA 92037; Omenn-Darling Bioengineering Institute, Princeton University, Princeton, NJ 08544

## Abstract

Protein interactions with DNA often result in rotational movements. These movements underlie many genetic processes such as DNA transcription by RNA polymerase (RNAP). To illuminate these movements, we previously developed Origami Rotor-Based Imaging and Tracking (ORBIT), a fluorescence-based method that enables high-speed tracking of DNA rotation. When used to track DNA rotation during transcription by *E. coli* RNAP, ORBIT enabled detection of single base-pair steps. However, the duration of ORBIT experiments was limited due to fluorescence photobleaching. To overcome this limitation and extend the total possible observation time, we here introduce dye-cycling ORBIT (dcORBIT), a method that enables observation of protein-DNA interactions for over 10 minutes at 20 Hz while maintaining single base-pair precision. dcORBIT thereby opens new possibilities to study the fundamental rotational movements underlying gene expression.

## Introduction

Many proteins cause rotation, bending, and other physical movements of DNA. Measuring these movements during DNA processing is critical for developing a full understanding of the mechanisms necessary for proper cellular function.

During transcription, RNAP moves processively along a DNA template as it synthesizes RNA. Due to the helical geometry of DNA, the translational movement of RNAP along the DNA is accompanied by a rotational movement of ∼35° per bp^1^. To track these rotational movements during transcription elongation, we developed Origami Rotor-Based Imaging and Tracking (ORBIT). In this method, stalled RNAP transcription elongation complexes are first attached to DNA origami rotors that each have a fluorescently labeled tip. These stalled RNAP-DNA-rotor complexes are then attached to a microscope cover glass via adsorption of RNAP to the glass surface. Transcription is then resumed, causing RNAP to pull and rotate the template DNA. The template DNA rotation can be followed via single-particle tracking of the labeled DNA origami rotor tip. Using this method, we directly observed ∼35° rotational steps during transcription elongation, corresponding to the helical twist between individual DNA base pairs^2^. ORBIT can therefore be used as a powerful method to study RNAP dynamics.

Similar to other fluorescence tracking methods, ORBIT observation times are limited by the photobleaching of the fluorescent dyes used to label the rotors. This photobleaching effectively limits the length of the DNA sequence over which we can track transcription at single bp resolution, and precludes the observation of RNAP dynamics over DNA stretches longer than a few base pairs. To study RNAP dynamics over longer time- and length scales without sacrificing base-pair resolution, it would be ideal to overcome this limit imposed by photobleaching.

Inspired by DNA-PAINT^3^, recent advances in fluorescent oligo dye-cycling has enabled single-particle tracking over longer time scales ^4–8^. In this dye-cycling strategy, the object to be tracked is tagged with a long single-stranded DNA oligo. This long oligo can then hybridize to multiple shorter fluorescent oligos (referred to as probes) present in solution. Keeping the hybridization length short causes the probes to only bind transiently before dissociating. By tuning the hybridization sequence, probe concentration in solution, and buffer conditions, one can ensure that at any given time there are multiple fluorescent probes bound to the tracked object. In this way, dye-cycling continuously replenishes the fluorescent probes and enables long-term single-particle tracking. We reasoned that application of dye-cycling in ORBIT would allow us to overcome photobleaching and track RNAP transcription at high angular and temporal resolution over long time scales.

Here, we introduce dye-cycling ORBIT (dcORBIT), a photobleach-resistant, single-molecule fluorescence imaging technique to follow dynamic protein-DNA interactions at base-pair resolution. First, we developed dye-cycling enabled DNA origami rotors compatible with ORBIT and identified optimal imaging conditions for continuous long-term tracking. Then, we validated dcORBIT by tracking DNA rotation during transcription with bp resolution over 10 minutes and measured the heterogeneity of RNAP step sizes during transcription elongation.

## Results

### Dye-cycling enables continuous ORBIT rotor imaging for 10 minutes

The DNA origami rotors consist of two perpendicular blades made of DNA helical bundles (Figure 1A), each containing six parallel DNA helices. In the original ORBIT method, each of the six DNA helices on the labeled tip were permanently labeled with a single fluorescent dye, yielding a total of six dyes per rotor. We hereafter refer to this configuration as “static-labeled”.

**Figure 1.**
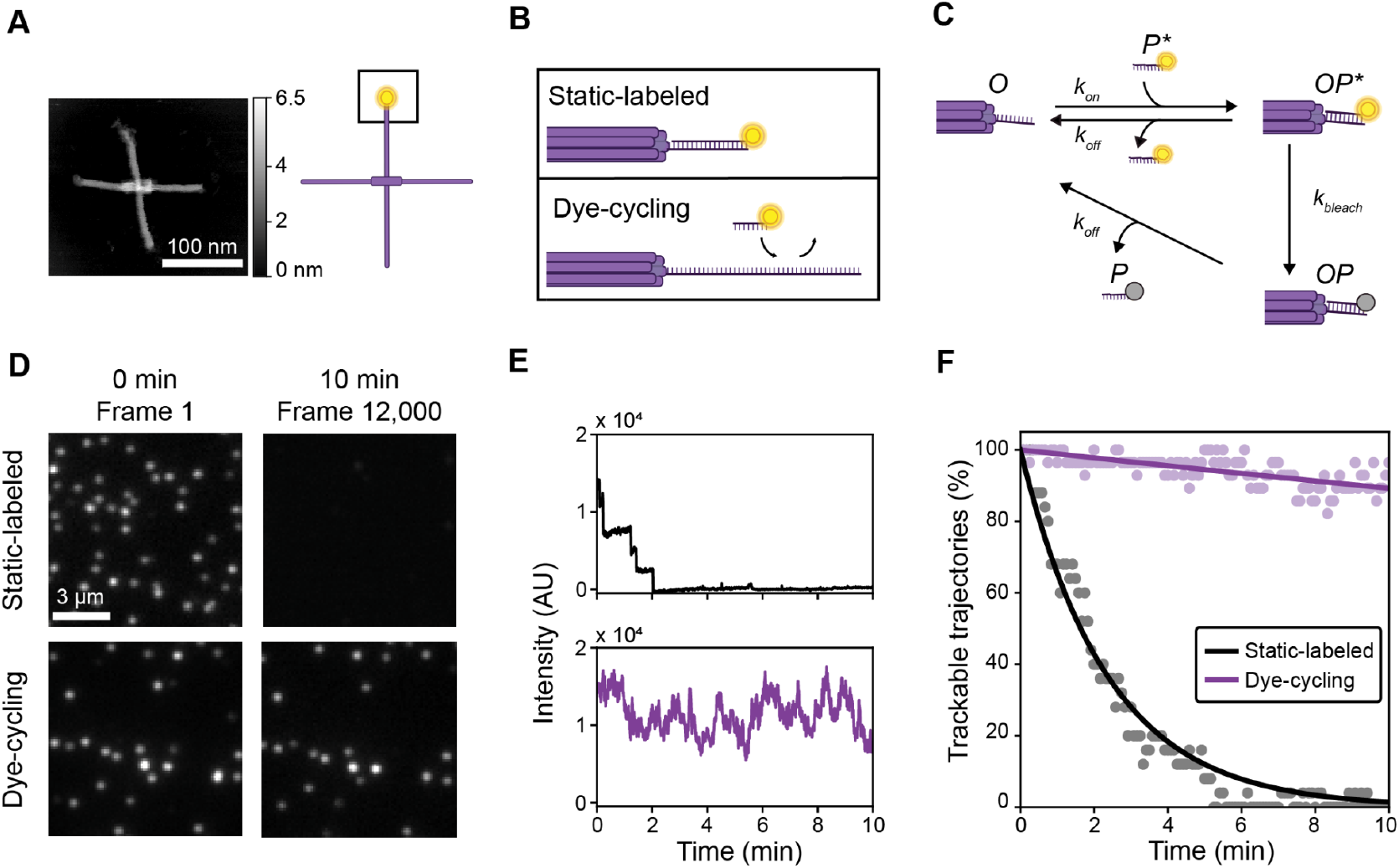
Dye cycling enables long-term observation of DNA origami rotors. **a** *Left*: Atomic force microscopy scan of a dye-cycling enabled DNA origami rotor. *Right*: Schematic drawing indicating location of fluorescent label on rotor blade tip. **b** Rotor blade tip close-up view showing static-labeled (*top*) and dye-cycling (*bottom*) configurations. The longer ssDNA overhangs on the dye-cycling rotors enable reversible binding of multiple 8-nt fluorescent oligos (probes) during imaging, thereby continuously replenishing the labels and avoiding the irreversible loss of fluorescence caused by photobleaching. *Note*: *Both static and dye-cycling configurations feature six ssDNA overhangs, one for each of the six DNA helices in the DNA origami rotor blades, however for clarity only one overhang in each configuration is shown here*. **c** Dye-cycling kinetic diagram. Overhangs (*O)* can reversibly bind fluorescent probes in solution (*P\**). Overhang-bound probes (*OP\**) can become photobleached (*OP*). Photobleached probes (*P*) can dissociate, leaving the overhang (*O)* open to bind a new probe (*P\**) in solution. **d** Total Internal Reflection Fluorescence (TIRF) microscopy of static-labeled rotors (*top row*) and dye-cycling rotors (*bottom row*). Left and right images in each row show the same field of view in frames 1 and 12,000, respectively. Static-labeled rotors exhibit near-complete photobleaching, while dye-cycling rotors maintain fluorescence throughout the recording. **e** Fluorescence intensity measurements of a representative static-labeled rotor (*top*) and a dye-cycling rotor (*bottom*). Data acquired at 20 Hz, background subtracted, and filtered using a 10-frame boxcar filter. **f** Percent of static-labeled rotors (*black*) and dye-cycling rotors (*purple*) in a field of view that retain sufficient fluorescence signal to enable base-pair (35°) precision tracking at 20 Hz.

To enable dye-cycling on the rotor, we added to each of the six helices a 44-nt single-stranded overhang with a sequence inspired by repetitive DNA-PAINT sequences, that included multiple overlapping binding sites for fluorescent probe oligos^4,5^ (Figure 1B, Figure S1). When imaging the rotor, we would then add a low concentration of a short dye-conjugated oligo probe to the imaging buffer. These short probes would transiently bind and dissociate from any of the multiple binding sites on the six overhangs during imaging, thus replenishing any photobleached probes. Importantly, our TIRF microscopy setup only illuminated the first few hundred nanometers above the cover glass surface, which minimized background fluorescence from dyes in solution and limited photobleaching of probes that were not near the surface.

To maximize the number of active fluorescent dyes on each overhang, we constructed a kinetic model, based on previous photobleaching optimization methods^9^, featuring the rates of probe binding (*k*_*on*_), unbinding (*k*_*off*_), and photobleaching (*k*_*bleach*_), as shown in Figure 1C. Importantly, we could tune these three rates by changing experimental parameters in a controlled manner. Specifically, we could change *k*_*on*_ through probe concentration, length, and sequence; *k*_*off*_ through probe length and sequence; and *k*_*bleach*_ through excitation laser intensity. By solving for the steady-state populations (Figure S3; see Methods), we could then select a combination of these rates that promoted continuous occupancy of multiple non-bleached probes on a rotor, under illumination conditions adequate for single-molecule tracking (Figure S4). We achieved these optimal rates by using a probe concentration of 20 nM and a probe length of 8 nt.

To test if dye cycling improved tracking, we compared the total amount of time that rotors could be tracked using static-labeled vs dye-cycling rotor tips. To do this, we imaged both kinds of rotors under the same image acquisition conditions for 10 minutes (Figure 1D). We chose these imaging conditions according to what we typically used to perform ORBIT experiments with static-labeled rotors. Initially, both kinds of rotors displayed a similar level of signal above the background. However, after 10 minutes, the static-labeled rotors had photobleached to near extinction, while the dye-cycling rotors retained their fluorescence throughout this time frame (Figure 1D). Fluorescence intensity measurements of a single representative static-labeled rotor showed successive photobleaching steps until the signal reached background levels (Figure 1E, *top*). Meanwhile, a representative dye-cycling rotor showed fluctuations of intensity likely arising from dynamic binding and unbinding of probes, but with a total signal that remained well above background levels throughout this measurement window (Figure 1E, *bottom*).

To test our ability to track the dye-cycling rotors throughout the 10-minute recording, we set a Signal-to-Noise Ratio (SNR) threshold based on the minimum SNR values that allow accurate rotational tracking of the rotor tip along the circular path (Figure S5A). The number of rotors above the SNR threshold in each labeling strategy was counted at every 100 frames and normalized to the total number of initially trackable rotors within that field of view (Figure 1F). Our results show that around 90% of the dye cycling rotors were still trackable after 10 minutes, while none of the static-labeled rotors were trackable after the same amount of time. Exponential fits to these data revealed that when compared to static-labeled rotors, dcORBIT enabled 37x longer observation times.

### Extended tracking of RNAP transcription with dye-cycling ORBIT rotors

ORBIT can be used to track DNA rotation during transcription by RNAP (Figure 2A). We had previously discovered that RNAP rotates the template DNA in discrete bp steps during transcription elongation^2^. To resolve these steps, we had to slow down transcription to a rate of ∼1 bp/s by lowering the free NTP concentrations, image with a frame rate of at least 20 frames per second, and illuminate the rotors with sufficient amount of excitation power to allow for nm-scale localization precision during these ∼50 ms exposure times. This combination of requirements led to our observations often being cut short by photobleaching, and as a result we rarely observed more than a few consecutive bp steps.

**Figure 2.**
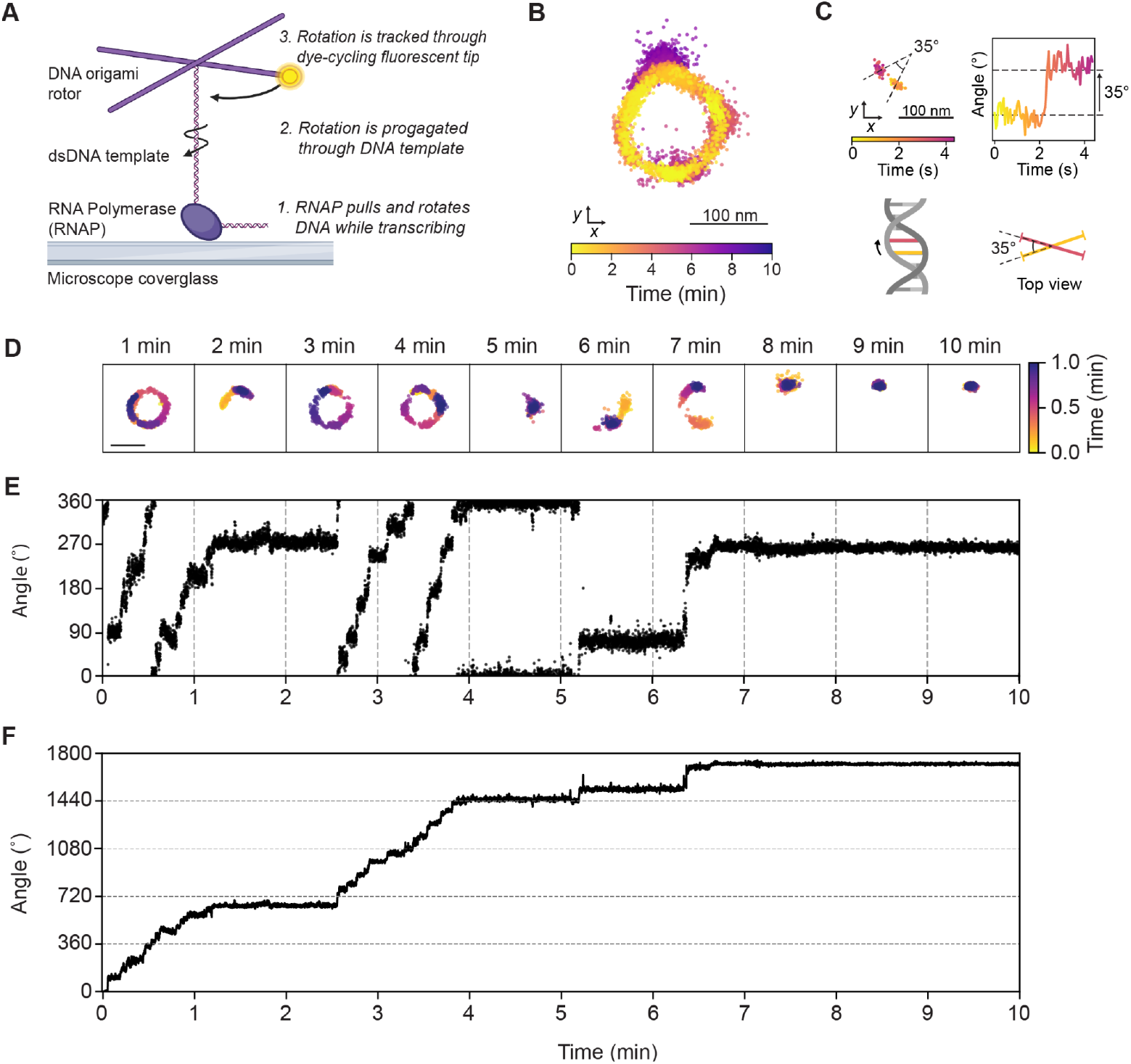
dcORBIT enables long-term tracking of RNAP transcription at base-pair resolution. **a** In the dcORBIT assay, surface-immobilized RNAP enzymes transcribe a DNA template attached to a DNA origami rotor. The processive movements of RNAP rotates the DNA template which drives rotor movement. Rotation is measured using single-molecule tracking of the dye-cycling DNA origami rotor tip. **b** Single-frame localizations of a rotor tip during RNAP transcription elongation show processive motion confined to a circular trajectory. Captured at 20 Hz over 12,000 frames at 10 µM NTPs. **c** dcORBIT resolves single base pair steps. *Top row*: Rotor tip localizations (*left*) and corresponding angle trace (*right*) showing a trajectory segment that contains a stepwise angular shift of ∼35^°^. *Bottom row:* The rotational twist between two adjacent base pairs in dsDNA is approximately 35°. **d** Rotor tip localizations from the trajectory in (*b*) separated into one-minute segments. Scale bar: 100 nm. **e** Rotor tip angle corresponding to localizations shown in (d). Rotational noise appears largely constant throughout trajectory, indicating that tracking precision is maintained over 10 minutes. Single base pair steps are visible throughout the first 4 minutes, followed by occasional larger movements consistent with RNAP reaching the end of the DNA template. **f** Cumulative rotation for the trajectory shown in (*e*). Horizontal dashed lines indicate intervals of 360^°^ which corresponds to ∼10.5 bp of transcription.

We conducted RNAP transcription experiments using the dcORBIT rotor, and a representative rotation tracking trajectory acquired at 20 Hz frame rate is shown in Figure 2B. The location of the rotor tip at each frame is represented by a point, with color indicating time. It is clear from this trajectory that the localization precision is sufficient to unambiguously determine the rotor’s angular position throughout the 10-minute long trajectory, as the rotor is being spun by a single RNAP enzyme. This trajectory contained clear examples of ∼35° steps, which corresponds to the expected rotation as the RNAP enzyme shifts the template DNA down and unwinds a single base pair (Figure 2C).

When displaying the trajectory from Figure 2B in 1-minute segments (Figure 2D) the RNAP-driven rotation can be seen to progress in spurts interspersed with pauses. Within each of these spurts, RNAP rotates the DNA in a series of single bp steps that occur at a relatively constant rate, which can be more easily seen in the corresponding angle trajectory (Figure 2E). Converting this trajectory to instead show the cumulative rotation of the DNA reveals more than four complete revolutions corresponding to more than 42 bp unwound (at 10.5 bp per revolution), with most of these steps being clearly resolved (Figure 2F). When compared to our previous study, dcORBIT thereby enables a nearly 10x improvement in our ability to detect consecutive bp steps during RNAP transcription.

### Heterogeneity of single-molecule RNAP transcription rates

At an NTP concentration of 10 µM (for each nucleotide), the transcription rates of individual RNAP enzymes varied greatly (Figure 3A). Assuming an angular shift of 35° per bp transcribed, these rates varied from 0.07 to 4.0 bp/s, with an average rate of 1.0 bp/s (Figure 3B). This average rate is in agreement with previously reported transcription rates for RNAP at 10 µM NTPs^1,10^, showing that the addition of dye-cycling in our assay did not significantly perturb RNAP transcriptional dynamics.

**Figure 3.**
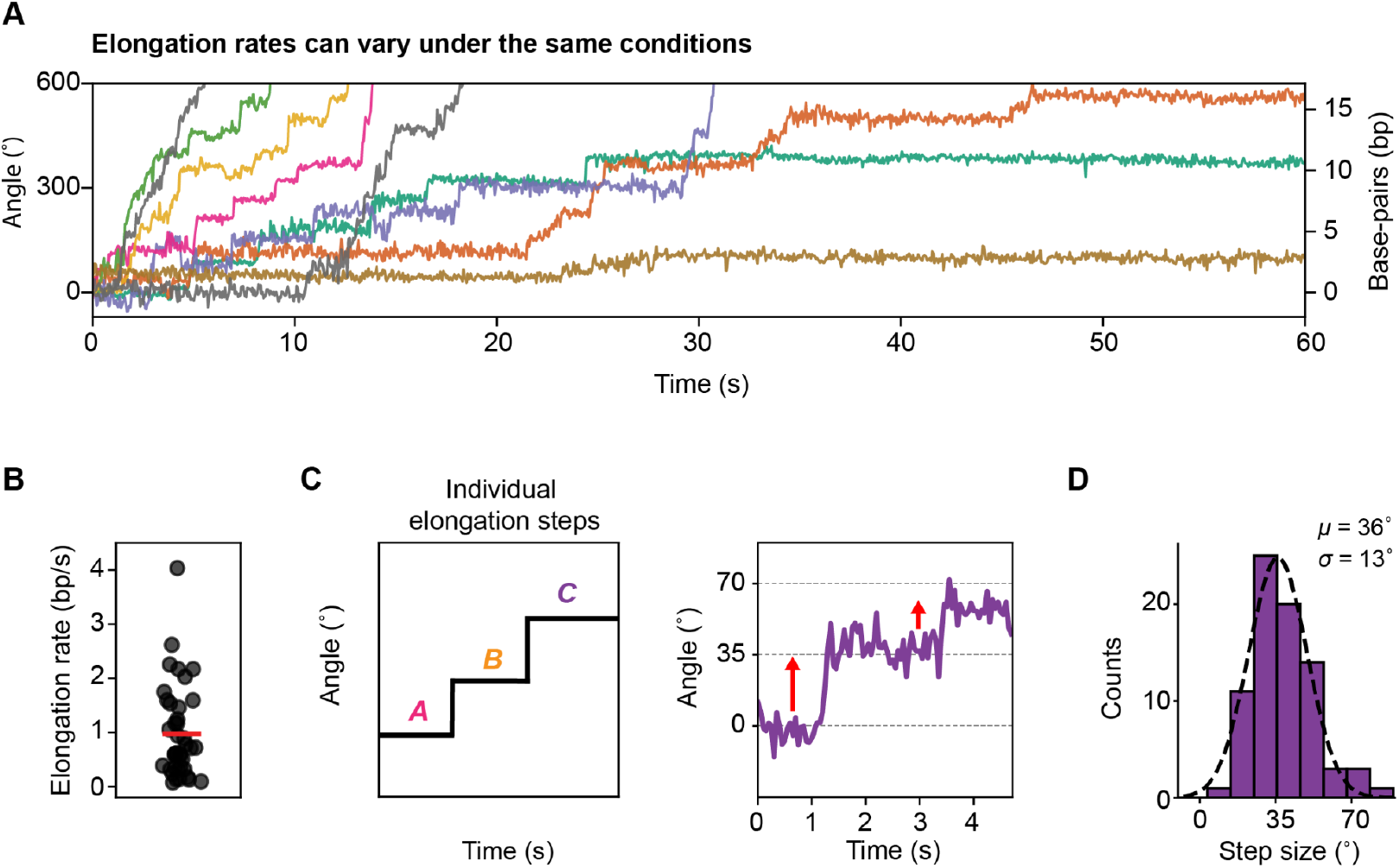
Heterogeneity of dynamics during transcription by RNAP. **a** Cumulative rotation vs. time for nine different individual RNAP enzymes show heterogeneity in rotational rates and stepping motion. **b** Single RNAP elongation rates at 10 µM NTPs. *N* = 43. Red line indicates the average rate of 1 bp/s. **c** Schematic and dcORBIT rotational trajectory (*purple*) showing processive forward base-pair stepping during RNAP transcription. **d** Histogram of forward step sizes during processive transcription. n = 78.

### Heterogeneity of rotational step sizes during RNAP transcription

Our previous study revealed that during transcription, the single bp steps by RNAP were not all equal in size^2^. To further examine this apparent heterogeneity of RNAP rotational step sizes, we measured the change in rotational angle during discrete processive forward steps across many single RNAP molecules (Figure 3C). The distribution of these forward step sizes was well captured by a Gaussian function centered at 36°, in agreement with the average helical twist of B-DNA, and with a standard deviation of 13.5° (Figure 3D). To assess whether the measured variation in step size could have been due to measurement error, we estimated our step size measurement uncertainty from the angle variance in the plateaus just before and after each step, and only included step measurements with an uncertainty less than 5°. Since the standard deviation of our measured step sizes was nearly threefold greater than the measurement uncertainty for any individual step (Figure S5B), we conclude that RNAP rotates the template DNA in elemental steps of uneven magnitudes during transcription.

## Discussion

ORBIT is a powerful method for the study of rotational movements during protein-DNA interactions. Here we describe the development of a photobleach-resistant version of ORBIT named dcORBIT, which enables more than an order of magnitude longer observation time window without compromising spatial or temporal resolution. Using dcORBIT, we tracked RNAP transcription at single-bp resolution for 10 minutes and were able to resolve the timing and magnitude of nearly every bp step along a transcribed template sequence.

The longer observation time windows with dcORBIT allowed us to measure the transcription rate of single RNAP enzymes at low nucleotide concentrations and in the absence of an external force. We found a substantial heterogeneity in the rates of individual enzymes, as these rates spanned nearly two orders of magnitude. Meanwhile, the ensemble average rate aligned well with previously reported values^1,10^, leading us to conclude that the enzyme-to-enzyme heterogeneity likely was not due to an artifact of our assay, but rather a reflection of previously observed differences between individual RNAP molecules that persist on a time scale of minutes^11,12^.

We further validated our previous discovery that the single-bp rotational steps by RNAP are of different magnitudes and measured the standard deviation in step sizes to be 13.5°.

With the development and validation of dcORBIT presented here we have made it possible to track transcriptional dynamics without the use of an external force, at bp resolution and over a time scale of several minutes. We envision dcORBIT to be a powerful assay for the study of a wide variety of protein-DNA interactions that induce rotation across a range of time scales such as other polymerases, helicases, remodeling enzymes, DNA repair enzymes, and gene editing enzymes^13–16^.

### Limitations of the study

The usage of a buffer containing fluorescent probes increases the fluorescence background of the experiment and necessitates the use of TIRF microscopy. This limitation could potentially be overcome by instead using fluorogenic probes^6^. The length of the template DNA on which we can track transcription is limited by the DNA persistence length (∼150 bp) and the height of the TIRF evanescent field (∼150 nm). We optimized our image conditions to achieve efficient dye-cycling given the imaging frame rate and localization precision required for single bp step detection during transcription; other applications of dcORBIT may require different imaging settings, necessitating a different set of optimal conditions for dye-cycling. This could be achieved using the optimization methods presented here.

## Methods

### DNA Origami Rotor Design

The static-labeled DNA origami rotors were made as described previously^2^, and the process is described in detail below in *DNA origami rotor folding, purification, and ligation*. Rotor blades are designed to be long enough to amplify the motion of the DNA template while maintaining low hydrodynamic drag and torsional flexibility. Each blade is formed from six DNA helices bundled together and rotors consist of four 80-nanometer (nm) “blades” that extend perpendicular to the axis of rotation. To enable static-label fluorescence tracking, one blade’s six DNA helices are each equipped with a 16-nucleotide (nt) single-stranded DNA overhang (Static-label overhangs in supplemental information) that is hybridized with a 16-nt fluorescently labeled probe (Static-label probe in supplemental information). To allow for linking to custom double-stranded DNA segments of experimental interest (Rotor linker strands in supplemental information), rotors contain a short dsDNA linker with a single-stranded overhang region at the center of where the four blades meet extending from the axis perpendicular to rotation.

### Dye-cycling rotor and probe design

Dye-cycling rotors have 44-nt overhangs extending from the same six DNA helices on one blade as static-labeled rotors (Dye cycling overhang sequences in Supplemental Table 1). The overhang sequence and complementary probe design are adaptations from a similar strategy used in Stehr et al.^5^ (Dye-cycling probe sequence in Supplemental Table1).

### DNA origami rotor folding, purification and ligation

All DNA origami staple strands, overhangs, linkers, fluorescently conjugated probes, and phosphorylated DNA were ordered from IDT with standard desalting or HPLC purification. DNA origami staples are generated using CADnano. DNA origami rotor staple strands (ORBIT staple strands in supplemental Table 1) were mixed equimolar, excluding overhangs, and linkers. DNA origami staple strands M13mp18 viral DNA purchased from New England Biolabs (N4040S).

DNA origami rotors were folded in a buffer containing 10 mM Tris, 1 mM EDTA, and 18 mM MgCl_2_. Folding occurred in a 50 µL total reaction with DNA at the following concentrations: M13mp18 scaffold at 10 nM, staple strands at 100 nM, DNA strands containing overhangs for fluorescent probes at 200 nM. Static-labeled rotors were folded with 500nM fluorescent DNA probes. Dye-cycling rotors were not folded with fluorescent DNA probes in the folding reaction. Reactions were incubated in a thermocycler and underwent a folding protocol starting at 80°C for minutes and then cooled to 65°C in 1°C steps every 5 minutes, followed by further cooling to 25°C in 1°C steps every 105 minutes. Folded rotors were then purified by PEG precipitation and stored frozen at 200 nM in TE buffer (10 mM Tris, 1 mM EDTA) + 18 mM MgCl_2_.

### Atomic force microscopy imaging

AFM images were acquired using the Dimension FastScan system (Bruker Corporation). Two µL of 1 nM sample was added to a freshly exfoliated mica disc. This was followed by a 20 µL droplet of buffer consisting of TE buffer, 9mM MgCl_2_, and 10mM NiCl_2_, allowed to incubate for 2 minutes, then another 150 µL of buffer was added to the mica disc. Imaging was done using C-type silicon tips (frequency, *f*0 = 56 kHz, spring constant, k = 0.24 N *·* m-1) on the nitride lever of the SNL-10 cantilever chip (Bruker Corporation).

### Static-label and dye-cycling kinetic controls using surface immobilized rotors

Static-labeled rotors and dye-cycling kinetics measurements were done on DNA origami rotors with a biotinylated linker strand allowing for surface immobilization. In both cases, cover glass (1.5, VWR 48404-467) was sonicated in 100% ethanol for seven minutes, dipped in MilliQ water, and pressure air dried. Flow chambers were assembled by adding double sided tape to ethanol cleaned glass microscope slides and placing the cleaned cover glass on top. After cover glass was added, double-sided tape is pressed using the back of a plastic tweezer to ensure proper adhesion and minimize liquid seeping through the tape. Flow chambers were then incubated with 10 mg/mL BSA-biotin (Thermo 29130) for 3 minutes, washed with TE buffer, incubated with 1 mg/mL Streptavidin (NEB N7021S) for 3 minutes, and then washed with TE buffer. Rotors with biotinylated linkers were diluted in TE buffer + 10 mM MgCl_2_, added to flow chamber and then incubated for 3 minutes before washing with TE buffer + 10 mM MgCl_2_. Standard imaging buffer of 250 µL containing 50 mM Tris, 0.5 mM EDTA, 10 mM MgCl_2_, 3 mM Trolox, 50 µM Trolox Quinone, 10% w/v glucose was prepared, and 5 µL of 42 mg/mL catalase and 1 µL of 50 mg/mL glucose oxidase is added fresh before flowing imaging buffer into flow chamber.

Static-label rotors had six overhangs statically labeled with a Cy5 probe. Static-labeled images were acquired at 20 Hz for 10 minutes. Videos were processed through the same data analysis pipeline as the transcription elongation videos.

Dye-cycling rotors used to find kinetic measurements were folded to have only one dye-cycling enabled overhang out of six (Dye-cycling kinetics overhang in supplemental information), and the overhang is shortened to 8-nt to enable capture of single binding and unbinding events of Cy5 probes in solution during imaging. The rotor tip opposite of the dye-cycling overhang had static-labeled overhangs with a complementary Cy3 probe to allow for localizing rotors independent of dye-cycling events (Rotor reference label and Reference label overhangs in Supplemental Table 1). An image was captured in the Cy3 reference channel to locate rotors in a field of view. Then, dye-cycling binding events were captured in the Cy5 channel for 1 minute at 2 Hz in an imaging buffer with 1 nM probes in solution at low laser power. In ImageJ, rotor locations were found in the Cy3 image and then used to highlight regions of interest in the Cy5 recording. The time that a Cy5 probe was bound or not bound at that rotor was used to calculate the time on and time off (*k*_*on*_ and *k*_*off*_).

### DNA template amplification

DNA template was PCR amplified (Q5 High-Fidelity 2X Master Mix, NEB M0492S), 500 nM of Forward Primer, 500 nM of Reverse Primer, 500 pM DNA template. The thermocycler was set to 98°C for 30 seconds, then for 37 cycles: 98°C for 10 seconds, 62°C for 15 seconds, 72°C for 10 seconds, followed by 70°C for 2 minutes. Next, amplified DNA was column cleaned to manufacturers directions (Zymo Clean and Concentrator - 25 Kit D4033) and eluted at 50 µL. Concentrated template DNA was digested overnight with XhoI (NEB R0146S) on a thermomixer set to 1000 rpm at 37°C. Then, DNA was column cleaned once again, eluted at 50 µL, and nanodropped to get exact concentration.

### RNAP stalling and ligation to rotors

E. coli RNA polymerase holoenzymes (NEB M0551S) were stalled on a DNA template of interest using as previously described for studying synchronized transcription elongation complexes^17^. In short, we used a 154 base-pair double-stranded DNA containing a T7A1 promoter sequence, followed by a 20 bp sequence with only 3 out of 4 bases to enable polymerase stalling (Annotated DNA template sequence available in supplemental information). RNAP (final concentration of 24 units/mL) was stalled with 10 nM DNA template, 25 nM photocleavable biotin-capture oligo, 250 μM ATP, GTP, and CTP, 100 μM of the dinucleotide ApU, in TX Buffer (20 mM Tris-HCL [pH 8.0], 50 mM KCl, 1 mM DTT, 0.1 mM EDTA [pH 8.0]), 10 mM MgCl_2_, 1% BSA for 20 minutes at 37°C on a thermomixer set to 1,000 RPM. Rifampicin was added for 5 minutes to remove improperly bound RNAP from the DNA template. The solution was then incubated with 8 ng streptavidin coated beads (ThermoFisher 65002) for 30 minutes at 20°C on a thermomixer set to 1,000 RPM. Beads were then washed to remove any uncaptured complexes. Beads were added to ligation reaction mixture (∼20 nM final concentration of DNA origami rotors, T4 Ligase (NEB M0202L, 6,400 units), in 1x T4 Ligase buffer) for 1.5 hours at 20°C on a thermomixer set to 1,000 RPM. Beads were washed to remove unligated rotors, and beads were transferred to TE buffer + 10mM MgCl_2_ + 20 mM NaCl to photocleave. Stalled complex beads were incubated in a 350 nm UV light box for 10 minutes to photocleave and remove stalled complexes from beads. Beads were then removed with a magnet. Stalled complexes used to acquire data within 3 - 4 hours of stalling.

### Dye-cycling RNA polymerase experiments

Cover glass (1.5, VWR 48404-467) was sonicated in 100% ethanol for 7 minutes, pressure air dried, sonicated in fresh 1 M KCl for 7 minutes, dipped in MilliQ water and pressure air dried. Double-sided tape was placed to create a flow chamber on standard glass microscope slides and cleaned cover glass was placed and pressed down onto the tape.

Dye-cycling rotors were ligated to stalled RNAP elongation complexes as described above. Chamber volumes were measured and washed with TE buffer + 10 mM MgCl_2_ + 20 mM NaCl and stalled complex reaction solution was added. Complexes were incubated for 2 minutes and then washed with 6 times chamber volume TE buffer + 10 mM MgCl_2_ + 20 mM NaCl + 1% BSA. Imaging buffer containing 20 nM Cy5 probes was supplemented with 0.8 µL of 43 mg/mL catalase and 4 µL of 50 mg/mL glucose oxidase in a low-bind Eppendorf tube and imaging buffer containing 20 nM Cy5 probes and 10 µM of all four NTPs, supplemented with catalase and glucose oxidase is prepared in another Eppendorf tube. Imaging buffer with probes but no NTPs was added to the flow chamber to check coverage and ensure dye-cycling rotors are in field of view. The microscope slide was removed, an imaging buffer with probes and NTPS was added to the flow chamber, the slide was returned to the microscope stage, and focus was reestablished. Videos are recorded at 20 Hz, for 10 minutes.

### Data analysis

Videos were analyzed using custom code written in Python (available on GitHub at github.com/pkosurilab/OMMxDORA). In brief, fluorescent rotors were identified by creating a median image of the first 300 frames, and the 300-frame median value was calculated for each pixel. Pixel intensity values over a set threshold and that were present for all 300 frames were selected as fluorescent peaks of interest. Peaks within 3 pixels of another peak were discarded. Pixel coordinates that pass these thresholds were highlighted as fluorescent peaks of interests and saved for full-video analysis. Peaks were centered in a 7 by 7-pixel region of interest (ROI), and a 2D Gaussian was fit over the pixels within the ROI in each frame of the video to collect the x-position, y-position of rotor blade, and integrated intensity. Signal and noise values calculated from the 2D Gaussian fit were also collected for each peak, for each frame. Traces were drift corrected using at least three stationary reference peaks from the same video. The median x and y locations of reference peaks were used to calculate global x- and y-shifts within the corresponding video. These values were then subtracted from the originally x- and y-positions calculated for every peak found in that video. Corrected x- and y-positions for the first 1000 frames were used to perform a least-squares fit to determine the geometric center of the circular trajectory. Rotational angles were determined by assigning localization positions from each frame an angle based on its position relative to the perimeter of the circle fit in the previous step.

Elongation rate in 3B was calculated by taking single RNAP transcription elongation traces that appeared to processively move for at least 600 degrees and using 35° to represent one base pair moved. Step-size measurements were determined by first marking two consecutive plateaus in rotational data that had at least 3 data points within each clear plateau. Medians of data points held between each plateau marked were registered as the rotational location of that step. Differences between the location of the two consecutive plateaus was used as an individual step size measurement. Standard error of measurement was also calculated, and only plateau measurements with less than 5 degrees of standard error of measurement were used in Figure 3D.

## Acknowledgements

AW was funded by National Institutes of Health T32 Training Grant (GM133351), the Ann Martinet Endowed Fund, the Jesse and Caryl Philips Award, the Mary K. Chapman Foundation Award, the Salk Women & Science Scientific Career and Professional Development Award, and the Dan and Martina Lewis Biophotonics Fellowship. This work was also supported by a Beckman Young Investigator Award (PK). We would like to thank Lauren Takiguchi who inspired us to ask if binding probes after DNA origami folding would be possible.

## Author contributions

A.W. and P.K. conceptualized the idea of applying a dye-cycling fluorescent strategy to DNA origami rotors. A.W. carried out planning, experimentation, and data analysis. B.J.T. developed RNAP stalling and rotor ligation protocols. R.J.F. adapted the kinetic model and contributed to data analysis. J.W. contributed to the data analysis pipeline. N.A.M. performed AFM imaging. A.W. and P.K. wrote the manuscript.

## Supplemental Information

Document S1 - Supplemental Figures S1 - S5

Supplemental Table 1 - DNA sequences

