## Supplemental Figures S1 - S5 for "Dye-cycling DNA origami rotors for long-term tracking of transcription at base-pair resolution"

**a**

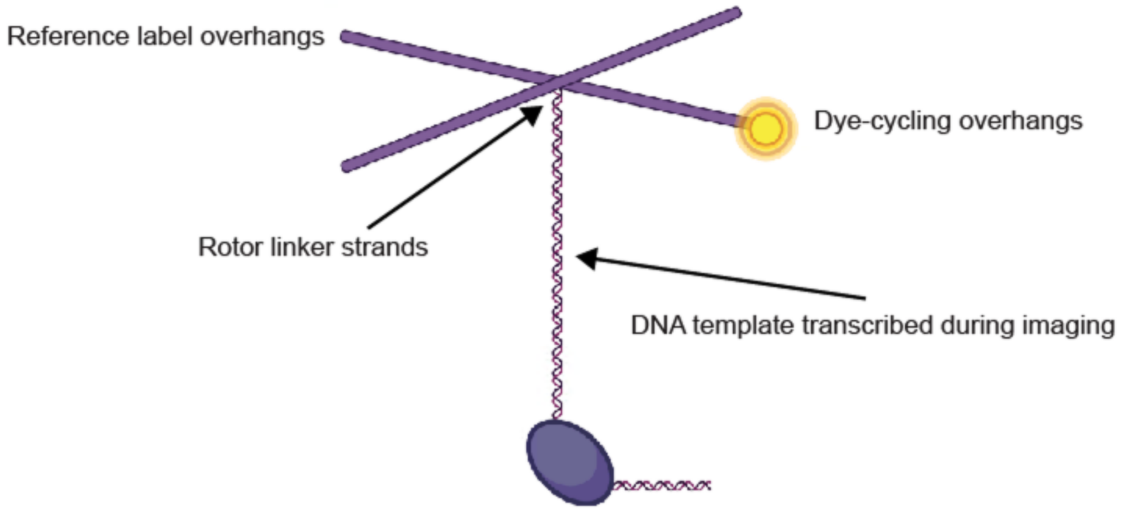


**Figure S1 | a** Schematic of ORBIT set up and where DNA sequences can be adjusted. Sequences included in additional information.

**a**

| Probe Name | Probe Sequence | $k_{on}$ [1/s • nM] | $k_{off}$ [1/s • nM] |
|---|---|---|---|
| 7A | 5cy5/AGGAGGA | 0.12 | 1.48 |
| 7G | 5cy5/GAGGAGG | 0.52 | 0.95 |
| 8 | 5cy5/GAGGAGGA | 0.02 | 0.03 |
| 9 | 5cy5/GAGGAGGAG | 0.001 | 0.01 |

**b**

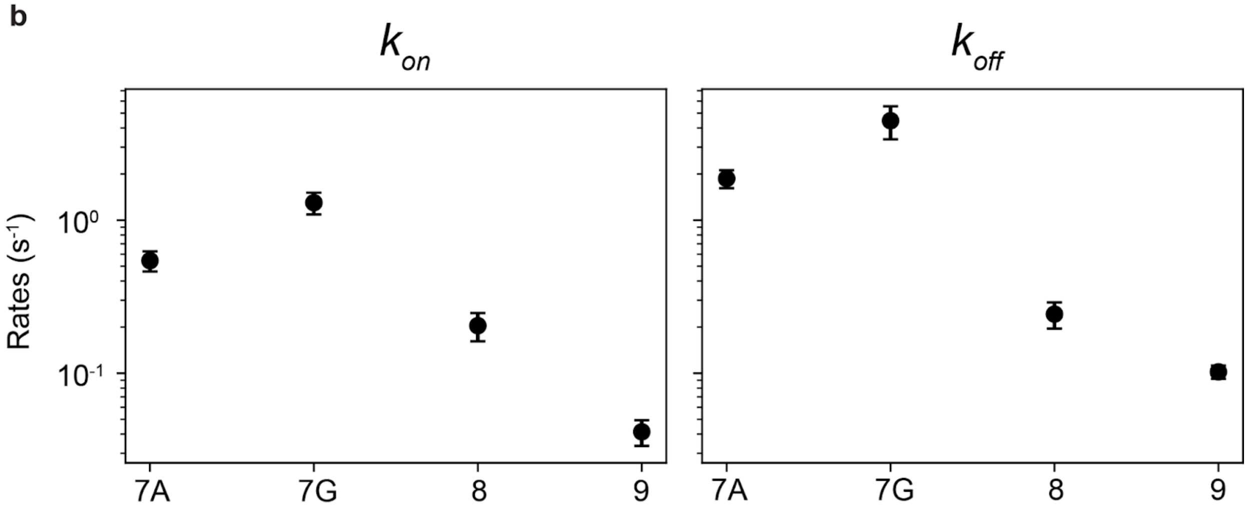


**Figure S2 | a** Tested dye-cycling probe sequences, experimentally measured rate constants. **b** Comparison of on-rate and off-rate of tested probe sequences at 1 nM probe concentration.

$$OP^* = \frac{1}{1 + \frac{k_{bleach} + k_{off}}{k_{on}} + \frac{k_{bleach}}{k_{off}}}$$

| **Parameters** | |
|---|---|
| **Description** | **Parameter** |
| On rate (1 nM P*) | $k_{on}$ |
| Off rate | $k_{off}$ |
| Bleach rate | $k_{bleach}$ |

| **Model** | | |
|---|---|---|
| **Description** | **Variable** | **Equation** |
| Overhang | *O(t)* | $\frac{dO}{dt} = k_{off}OP^* + k_{off}OP - k_{on}O$ |
| Overhang w/ Probe | *OP*(t)* | $\frac{dOP^*}{dt} = -k_{off}OP^* + -k_{bleach}OP^* + k_{on}O$ |
| Overhang w/ Bleached Probe | *OP(t)* | $\frac{dOP}{dt} = -k_{off}OP + k_{bleach}OP^*$ |

| **Assumptions** | |
|---|---|
| **Description** | **Assumption** |
| Concentration of probes is large | $\frac{dP^*}{dt} = 0, k_{on} = kP^*$ |
| Total binding sites are conserved | $O_{total} = O + OP^* + OP$ |

**Figure S3 | a** Dye-cycling model equation used to optimize probability of an overhang being bound by a probe that is able to emit fluorescence.

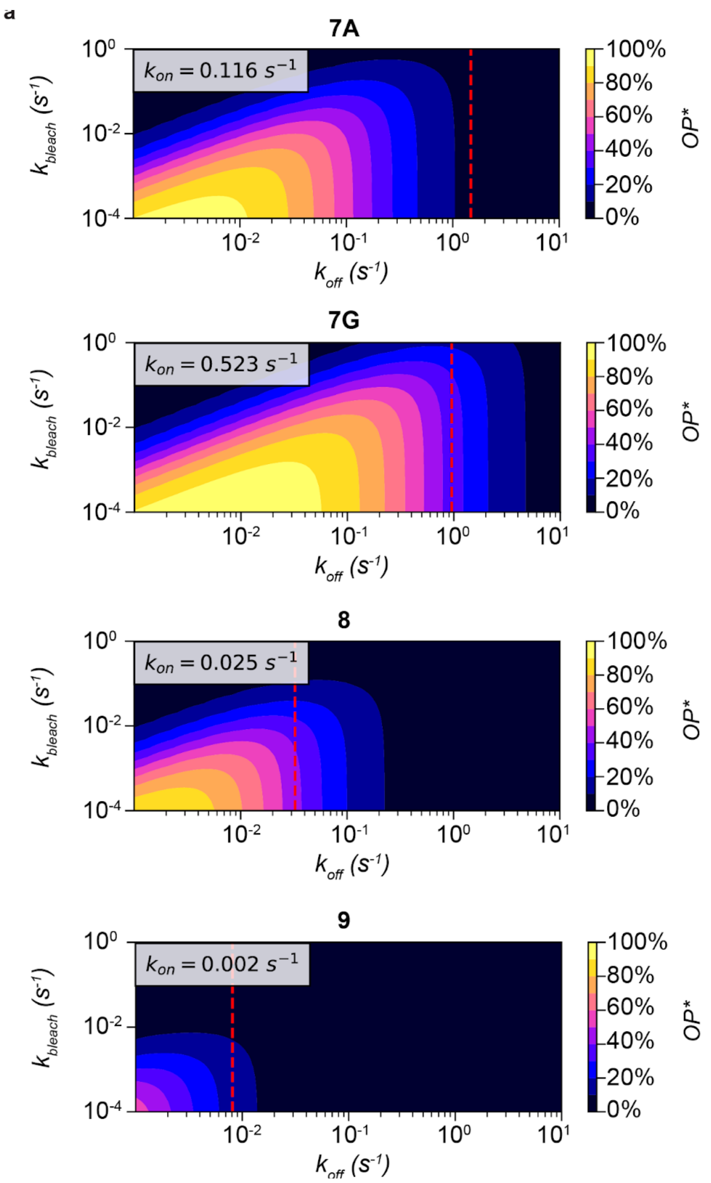


**Figure S4 |** Probability landscapes using rates found in Supplemental Figure S2 and dye cycling kinetic model described in Supplemental Figure S3.

**a**

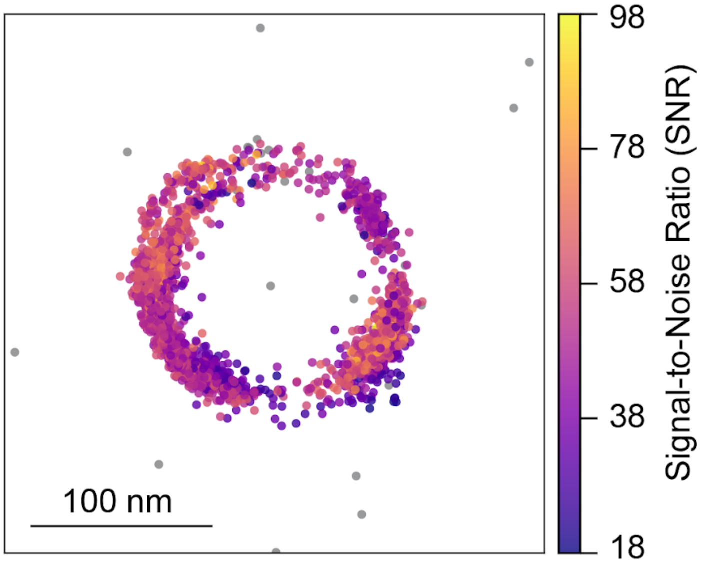


**b**

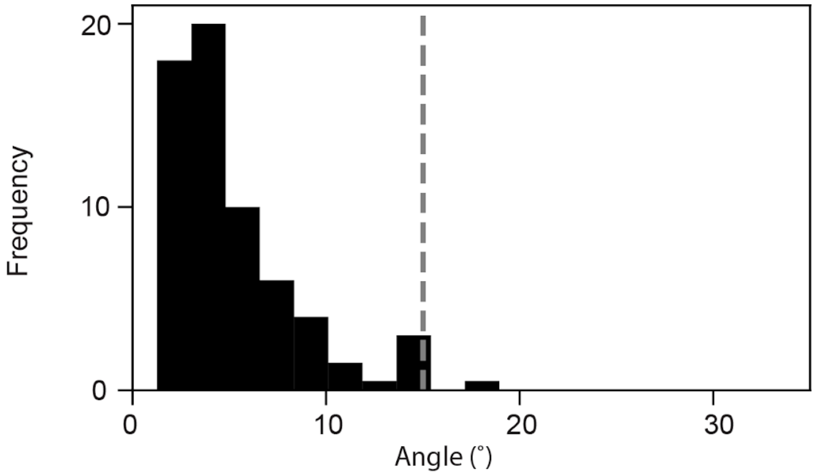


**Figure S5 | a** Example rotor localization during dye-cycling recording colored based on Signal to Noise Ratio (SNR) threshold. Observations with SNR less than 18 colored in grey. **b** Histogram standard error of measurement of unfiltered forward step size measurements. Gray vertical line indicates standard deviation of forward step sizes.
